# Identification of a novel, NF-κB nucleolar stress response pathway

**DOI:** 10.1101/100255

**Authors:** Jingyu Chen, Ian T Lobb, Pierre Morin, Sonia M Novo, James Simpson, Kathrin Kennerknecht, Fiona Oakley, Lesley A. Stark

## Abstract

p53 as an effector of nucleolar stress is well defined, but p53 independent mechanisms are largely unknown. Like p53, the NF-κB transcription factor plays a critical role in maintaining cellular homeostasis under stress. Many stresses that stimulate NF-κB also disrupt nucleoli. However, the link between nucleolar function and activation of the NF-κB pathway is as yet unknown. Here we demonstrate that siRNA silencing of PolI complex components stimulates NF-κB signalling. Unlike p53 nucleolar stress response, this effect does not appear to be linked to inhibition of rDNA transcription. We show that specific stress stimuli of NF-κB induce degradation of a critical component of the PolI complex, TIF-IA. This degradation precedes activation of the NF-κB pathway and is associated with an atypical nucleolar architecture. It is mimicked by CDK4 inhibition and is dependent upon upstream binding factor (UBF) and p14ARF. We show that blocking stress effects on TIF-IA blocks their ability to activate the NF-κB pathway. Finally, using *ex vivo* culture, we show a strong correlation between degradation of TIF-IA and activation of NF-κB in freshly resected, human colorectal tumours exposed to the chemopreventative agent, aspirin. Together, our study provides compelling evidence for a new, NF-κB nucleolar stress response pathway that has in vivo relevance and therapeutic implications.

## Introduction

The nucleolus is a highly dynamic, sub-nuclear organelle that, in addition to its primary function as the hub of ribosome biogenesis, acts as a critical stress sensor and coordinator of stress response (1,2). The starting point of ribosome biogenesis is transcription of ribosomal DNA (rDNA), which is mediated by the RNA polymerase I (PolI) complex (3). If cellular homeostasis is challenged, a variety of kinases target this complex and consequently, rDNA transcription is inhibited, the gross architecture of the nucleolus is altered, the nucleolar proteome is dramatically modified and ultimately, signals are transmitted to downstream effector pathways so that cell growth and death are altered accordingly (1,4,5). This chain of events is broadly termed “nucleolar stress” and the most recognised and characterised effector pathway is the MDM2-p53 axis (6-8). However, it is increasingly apparent that nucleolar stress can regulate cell phenotype in a p53 independent manner (9-13). Indeed, the mechanisms that coordinate stress effects on the PolI complex, and integrate these into individual phenotypic outcomes, remain poorly understood.

Similar to p53, the NF-κB transcription factor plays a critical role in regulating cell growth and death in response to stress (14). The most abundant form of NF-κB is a heterodimer of the p50 and RelA (p65) polypeptides which is generally bound in the cytoplasm by the inhibitor, IκBα (15). Upon exposure of the cell to a myriad of stresses, IκBα is degraded allowing NF-κB to translocate to the nucleus where it regulates expression of target genes (16-18). In contrast to the rapid activation observed in response to classical NF-κB stimuli, stress stimuli (including serum starvation, UV-C radiation and chemopreventative/therapeutic agents) generally induce this pathway with a much slower and delayed kinetic (19). Although a number of mechanisms have been proposed, how multiple environmental and cytotoxic stimuli induce the delayed activation of NF-κB remains unclear.

In this lab, we noted that a common response to stress stimuli of the NF-κB pathway is modulation of nucleolar architecture. In particular, an increase in the size of the organelle (19). This was of interest because, a common denominator of stresses that activate NF-κB is inhibition of rDNA transcription (some of which are summarised in Supplemental Table 1)(20-22). Furthermore, proteins that have a role in stress-mediated activation of NF-κB reside within this organelle (23-26). We have previously demonstrated that post induction, RelA can accumulate in nucleoli (19,27) and others have shown modulation of NF-κB signalling by ribosomal proteins (28,29). However, to the best of our knowledge, no association between nucleolar stress and induction of NF-κB signalling has previously been reported.

Here we investigated the relationship between stress effects on the PolI complex and NF-κB signalling. Our results reveal a novel nucleolar stress response pathway that culminates in activation of NF-κB. The pathway is characterised by CDK4 inhibition-UBF/p14ARF-mediated degradation of the PolI complex component, TIF-IA. It is triggered in a whole tissue setting and has relevance to the anti-tumour effects of aspirin.

## Materials and Methods

### Cell lines and treatments

Human SW480, HRT18, RKO and HCT116 colon cancer cells, PNT pancreatic cells and Hela Cervical cancer cells, are available from the American Type Culture Collection (ATCC). The p53 null derivative of HCT116 (HCT116^p53-/-^) was a gift from Professor B Vogelstein (John Hopkins University School of Medicine, USA) and has previously been described(30). HRT18SR, a derivative of HRT18 cells that constitutively expresses a non-degradable IκBα, was generated in this lab and has been described(31). All cells lines were maintained at 5% CO2 in growth medium (Gibco) supplemented with 10% fetal calf serum (FCS) and 1% penicillin/streptomycin. Medium used was: SW480: Leibovitz’s L-15: PNT, RKO, HCT116, HCT116p^53-/-^-DMEM; HRT 18, HRT18SR-RPMI with Geneticin (Gibco) selection.

All treatments were carried out in reduced serum (0.5% FCS) medium for the times and concentrations specified. Aspirin (Sigma) was prepared as previously described(31). ActinomycinD (Sigma), Cyclohexamide (Sigma), TNF (R&D Systems), ceramide C2/C6 (Sigma), MG132 (Sigma), lactacystin (Calbiochem), and quinacrine (Sigma) were all prepared as per manufacturer’s instructions and used at the concentrations given. For UV-C treatments, cells were mock treated or exposed to UV-C under the conditions stated. The CDK4 inhibitors 2-bromo-12,13-dihydro-5*H*-indolo[2,3-*a*]pyrrolo[3,4-*c*]carbazole-5,7(6*H*)-dione (CDK4i, Calbiochem) and Palbociclib (PD-0332991, Selleckchem) were solubilised in DMSO and used as indicated. The rDNA transcription inhibitor BMH-21 (12H-Benzo[g]pyrido[2,1-b]quinazoline-4-carboxamide, N-[2(dimethylamino)ethyl]-12-oxo) was kindly supplied by Prof. Marikki Laiho (Johns Hopkins University School of Medicine, USA) and Nutlin-3 by Prof. Kathryn Ball (University of Edinburgh, Edinburgh Cancer Research Centre, UK).

### Immunocytochemistry, image quantification and FUrd assays

Immunocytochemistry was performed as previously described (27). Primary antibodies were TIF-IA (BioAssayTech), RelA (C-20), Nucelolin (MS-3), RPA194 (all Santa Cruz Biotechnology) and Fibrillarin (Cytoskeleton). Cells were mounted in Vectashield (Vector Laboratories) containing 1ug/ml DAPI. Images were captured using a Coolsnap HQ CCD camera (Photometrics Ltd, Tuscon, AZ, USA) Zeiss Axioplan II fluorescent microscope, 63 × Plan Neofluor objective, a 100 W Hg source (Carl Zeiss, Welwyn Garden City, UK) and Chroma 83 000 triple band pass filter set (Chroma Technology, Bellows Falls, UT, USA). Image capture was performed using scripts written for IPLab Spectrum 3.6 or iVision 3.6 in house. For each experiment, a constant exposure time was used. Image quantification was carried out using DAPI as a nuclear marker and fibrillarin as a nucleolar marker along with ScanR (Olympus), IPLab or ImageJ image analysis software as specified in figure legends. At least 150 cells from at least 5 random fields of view were quantified for three independent experiments, or as specified in the text.

For fluorouridine (FUrd) run on assays, cells were treated with 2mM FUrd 15min before harvest. Immunocytochemistry was then performed with an anti-BrdU antibody (Sigma). Images were captured using a ScanR high-content imager (Olympus) with a LUCPLFLN 40X objective (Olympus) and ScanR Acquisition software (Olympus). FUrd incorporation was quantified for at least 1000 cells per slide using ScanR analysis software with particle recognition algorithms.

### Plasmids, siRNA and transfections

Flag-UBF wild type, S388G and S484A mutants were kindly provided by R. Voit (German Cancer Research Centre, Heilderberg, Germany)(32). NF-κB reporter constructs (3 x enhancer κB ConA (3x κB ConA-Luc), IκBα luciferase (IκBα-Luc)) and ΔB-deleted derivatives (ΔκB ConA-Luc, ΔIκBα-Luc) were provided by RT Hay (University of Dundee, Dundee, UK) and have been described elsewhere(19). pCMV-β is commercially available (Promega). pEGFP-C1-hTIF-IA was kindly gifted by I Grummt (German Cancer Research Centre, Heilderberg, Germany). Flag-P14ARF was gifted by A Lamond (University of Dundee, Dundee, UK). siRNA duplex oligonucleotides were synthesized by MWG and transfected into cells using lipofectamine 2000 following the manufacturer’s instructions. Cells were transfected on two consecutive days then left to recover for 24-48h prior to treatment or harvest. siRNA sequences are as follows: TIF-IA CUAUGUAGAUGGUAAGGUU; UBF CCAAGAUUCUGUCCAAGAA; p14ARF AAGACCAGGUCAUGAUGAUGG; Control AGGUAGUGUAAUCGCCUUG.

### Quantitative PCR

RNA was extracted from cells using RNeasy mini kit (Qiagen) following the manufacturer’s instructions. Extracted RNA was purified using RQ1 RNase-free DNase (Promega) then cDNA generated using 1st Strand cDNA synthesis kit (Roche). Taqman assays (Thermo Fisher Scientific) and a LightCycler 480 system were used to quantify transcript levels. Primers and probes used in taqman gene expression assay are given in full in additional information. The Comparative C_T_ Method (or ΔΔC_T_ Method) was used for calculation of relative gene expression.

### Immunoblotting, luciferase reporter and apoptosis assays

Immunoblotting, luciferase reporter and AnnexinV apoptosis assays were carried out as described previously (19,31). Primary antibodies used for immunoblots are as follows: TIF-IA (rabbit, 1:2000, BioAssayTech B8433); Rrn3 (Mouse, 1:500, Santa Cruz sc-390464); RPA194 (POLR1A) (H 300, Rabbit, Santa Cruz, sc-28714); RelA^S536^ (Rabbit, 1:500, Cell signalling, 3031S); GFP (Rabbit, 1:1000 Santa Cruz, sc-8433) UBF (Mouse, 1:500, Santa Cruz, sc-13125); UBF^S484^ (Mouse. Assaybiotech, A8444); p53 (Mouse, 1:2000; Oncogene OP43) IκB (Sheep, 1:5000, gift from RT Hay, University of Dundee).

### Immunoprecipitation

Immunoprecipitation assays were performed using 1mg whole cell lysate, prepared in NP40 lysis buffer. Mouse TIF-IA antibody (Santa Cruz Biotechnology) was used to immunoprecipitate the appropriate protein. Mouse IgG (pre-immune serum) was used as a control. Complexes were resolved by SDS polyacrylamide gel electrophoresis then analysed by western blot analysis.

### Ex vivo treatment of tumour biopsies and immunohistochemistry

Biopsies of colorectal tumours were provided by a pathologist at the time of resection. All patients were consented and full ethical approval was in place (Scottish Colorectal Cancer Genetic Susceptibility Study 3; Reference: 11/SS/0109). Biopsies were immediately transferred to the lab immersed in culturing media (MEM supplemented with glutamine, penicillin/streptomycin and anti-mycotic/antibiotic mix (1:100, Sigma). Tumours were washed, dissected into 1-2mM fragments then plated. Treatment (0-100μM aspirin, 1h, 37°C) of tumour explants was performed in 96-well plates in duplicate in the presence of 10% foetal calf serum (33). Following treatment, tumours were either frozen for protein analysis (set 1) or formalin fixed for immunohistochemistry (set 2). Whole cell extracts were prepared using a TissueLyser(Qiagen) and standard whole cell lysis buffer.

Anti-^p536^RelA immunohistochemistry was carried out using a DAB protocol on formalin fixed sections, as previously described (34). A Leica scanner digitised images then Leica QWin plus image analysis software (Leica Microsystems Inc., Buffalo, IL) used to analyse cells for nuclear RelA^p536^ staining. Three distinct areas of tissue and at least 1500 cells were analysed per section (average 4650). The quantification software determined the percentage of cells showing negative, weak, moderate and strong RelA^p536^ staining.

## Results

### Silencing of PolI complex components activates the NF-κB pathway

To investigate the link between nucleolar perturbation and NF-κB, we firstly inactivated an essential component of the PolI pre-initiation complex, upstream binding factor (UBF), then examined NF-κB pathway activity. This mimicked the approach used by Rubbi and Milner to show nucleolar stress stabilises p53(7). We found siRNA silencing of UBF caused an increase in S536 phosphorylated RelA(a marker for cytoplasmic activation of the NF-κB pathway), a significant increase in nuclear RelA and an increase in NF-κB-driven transcription comparable to that observed when cells were treated with the classic NF-κB stimulus, TNF (Figure 1*A*). Degradation of IκBα, pRelA^S536^ phophorylation and increased NF-κB-driven transcription were also observed upon silencing of the PolI components, POLR1A (RPA194) and TIF-IA (Rrn3p) (Figures 1*A* and B).

**Figure 1:**
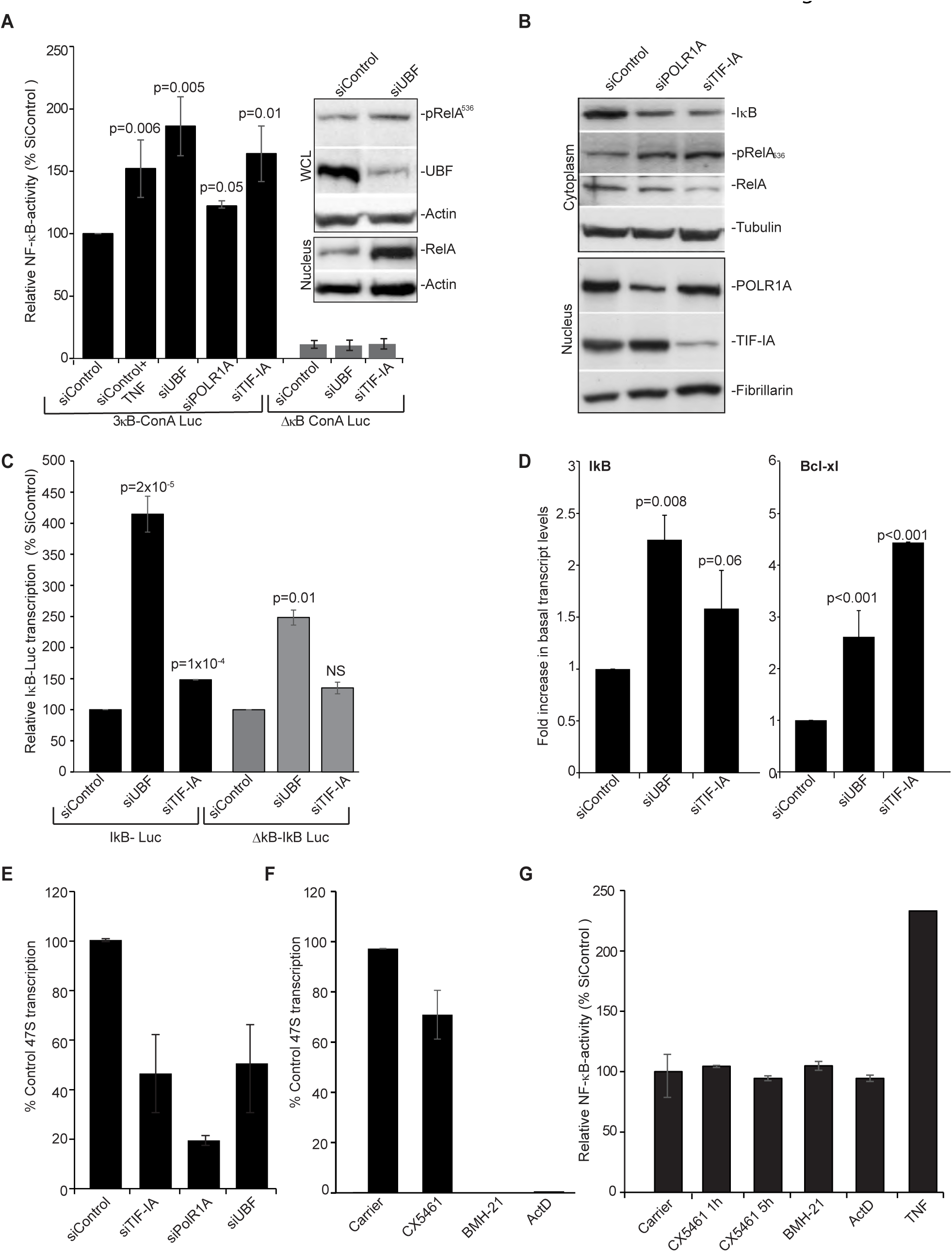
Silencing PolI complex components stimulates the NF-κB pathway. **(A-E)** SW480 (or **(D)** HCT116) cells were transfected with the indicated siRNA species **(A)** Cells were co-transfected with pCMVβ and either a wild-type NF-κB-dependent luciferase reporter construct (3x κB ConA) or an equivalent plasmid with κB sites deleted (ConAΔκB). TNF (10ng/ml, 4h) acts as a control NF-κB stimulant. Luciferase activity was normalized using β-galactosidase activity. Results are presented as the percentage of relative NF-κB activity compared to control siRNA (siControl). The mean of at least three individual repeats (±s.e.m) is shown. Inset: Levels of pRelA^536^, nuclear RelA and UBF in whole cell (WCL) or nuclear lysates were determined upon UBF silencing using Western blot analysis. Actin acts as a control. **(B)** Immunoblot demonstrates reduced cytoplasmic IκBα, increased pRelA^S536^ and decreased cytoplasmic RelA upon silencing of TIF-IA and POLR1A. Nuclear extracts confirm efficient protein depletion. α-Tubulin and fibrillarin act as loading controls. **(C)** Cells were co-transfected with IκB-luc (luciferase driven by full length IκB promoter) or ΔκB-IκB-luc (equivalent in which NF-κB sites are deleted) and pCMVβ. The percentage relative luciferase activity compared to siControl was calculated. Mean± s.e.m is shown (N=3). **(D)** qRT-PCR with primers for the NF-κB target genes IκBα and Bcl-xl. GAPDH was used to normalise. Results are presented as the fold increase in transcript compared to siControl. The mean (± s.e.m) is shown. N=3. **(E)** qRT-PCR with primers for the 47S pre-rRNA transcript measured levels of rRNA transcription. GAPDH was used to normalise. Results are presented as the percentage of relative 47S transcription compared to siControl. The mean (± s.e.m) is shown. N>3. **(F)** SW480 cells were treated with the PolI inhibitors CX5461 (500nM), BMH-21 (4uM), ActinomycinD (ActD, 1ug/ml) or TNF (10ng/ml), for 5h. qRT-PCR measured levels of the 47S transcript as above. Mean ± s.e.m is shown (N=2). (G) SW480 cells were transfected with 3x κB ConA-Luc and pCMVβ. Twenty-fours hours later they were treated with inhibitors as in F. Graph shows the mean of at least two individual repeats ±s.e.m. *P* values throughout are compared to the respective control and were derived using a two tailed students T test. See Supplemental Figure 1 for additional cell lines and supporting data.

A link between PolI complex disruption and increased NF-κB activity was confirmed using independent cell lines (Supplemental figures 1*A* and *C*). It was also confirmed using an independent reporter plasmid in which transcription of the luciferase gene is driven by the full-length promoter of the classic NF-κB target, IκBα (Figure 1*C*). Indeed, depletion of UBF caused a >5 fold increase in transcription from the IκBα promotor. However, this was significantly abrogated when an equivalent reporter plasmid lacking κB sites was utilised indicating NF-κB dependency. qRT-PCR confirmed that silencing PolI complex components induces transcription of IκBα and an independent NF-κB target, Bcl-xl (Figure 1*D*).

Given that disrupting nucleoli is linked with dramatic changes in the nucleoplasmic proteome^10^, we did consider that silencing PolI complex components may stimulate NF-κB-driven transcription without degradation of IκB/cytoplasmic release of NF-κB. However, the significant increase in NF-κB-driven transcription observed in control cells upon depletion of UBF and TIF-IA was blocked in cells we generated to constitutively express super repressor (non-degradable) IκBα (Supplemental figure 1*C*). Hence, we conclude that IκBα degradation is an essential step for nucleolar stress to induce NF-κB transcriptional activity.

qRT-PCR for the 47S pre rRNA transcript confirmed that siRNA to UBF, POLRIA and TIF-IA inhibited rDNA transcription (Figure 1*E*). Stabilisation of p53 is strongly linked to inhibition of rDNA transcription (8). Therefore, we considered this may also be the case for activation of NF-κB. However, we found that actinomycinD and two highly specific small molecule inhibitors of PolI (CX5461 and BHM-21(35,36)), caused a significant reduction in levels of the 47S transcript, but had no effect on NF-κB transcriptional activity (Figures 1*F* and *G*).

Taken together, these data suggest that it is not inhibition of rRNA transcription *per se*, but a specific type of perturbation of the PolI complex that activates the cytoplasmic NF-κB pathway. To test this suggestion, we next examined the effects of NF-κB stress stimuli on this complex.

### Degradation of TIF-IA precedes NF-κB pathway activation

TIF-IA is essential for rDNA transcription as it tethers Pol I to the rDNA promoter(37,38). It is also *the* component of the pre-initiation complex that is targeted by stress/environmental signals to alter PolI activity(39,40). Therefore, we firstly investigated this protein. Aspirin was initially chosen as a model stimulus as we are interested in the pro-apoptotic activity of this agent and, in the absence of additional cytokines, it stimulates NF-κB in a manner characteristic of multiple stress inducers(19,31).

Figure 2A demonstrates that aspirin not only induces a decrease in Ser649 phosphorylated TIF-IA, which is a known response to environmental stress, but also a significant decrease in native TIF-IA, which is *not* a reported stress response (Figure 2*A*). This decrease in native TIF-IA was observed by Western blot analysis (Figure 2*A*) and by immunocytochemistry (Figure 2*B*). It was evident in multiple cell types, was independent of p53 status and most importantly, was an early response to the agent preceding degradation of IκB and nuclear translocation of RelA (Figures 2*C* and *D* and Supplementary Figures 2A and B).

**Figure 2.**
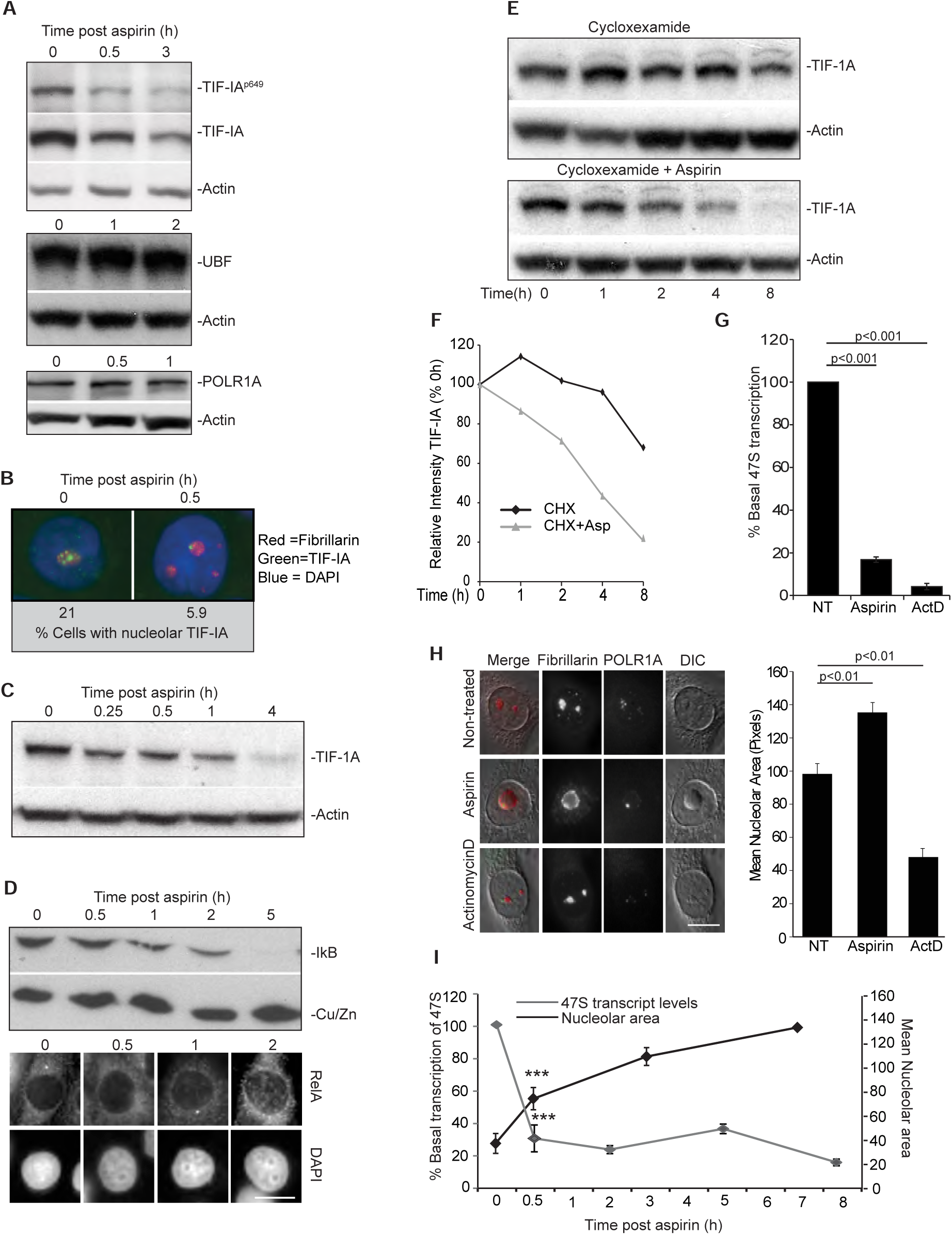
Degradation of TIF-IA and a distinct nucleolar phenotype precede aspirin effects on the NF-κB pathway. **(A to D)** Aspirin (used as a model stress stimuli) induces a decrease in total cellular levels of TIF-IA, which precedes degradation of IκB and nuclear accumulation of RelA. SW480 cells were treated with aspirin (10mM) for the indicated times **(A, C and D)** Immunoblot analysis was performed on WCL with the indicated antibodies. **(B)** Immunomicrographs (63X) show levels and localisation of native TIF-IA. Below: The percentage of nuclei (as depicted by DAPI stain) with bright puncti of TIF-IA were quantified using ImageJ software. A minimum of 200 nuclei were analysed per experiment from at least 10 fields of view. N=3. **(D)** Bottom: Immunomicrographs (X63) demonstrate accumulation of RelA in nuclei 2h after aspirin exposure. **(E and F)** SW480 cells were treated with cyclohexamide (10uM) alone or with aspirin (10mM) for the times specified. **(E)** Immunoblot shows cellular levels of TIF-IA. **(F)** TIF-IA band intensities relative to actin were quantified using ImageJ. Results are presented as the percentage compared to the 0h control. N=2. **(G to I)** Perturbation of nucleolar structure and function in response to aspirin. **(G)** rDNA transcription was quantified after aspirin (3mM, 16h) or actinocycinD (50ng/ml, 2h) treatment using qRT-PCR for the 47S transcript as above. Results are presented as the percentage of relative transcription compared to non-treated (NT) control. The mean (± s.e.m) is shown. N=4. **(H)** Representative DIC immunomicrographs (x63) showing the cellular localisation of components of the tripartite nucleolar structure in response to aspirin (10mM, 8h) and actinomycinD (50ng/ml). Fibrillarin marks the dense fibrillar component and POLR1A the PolI complex in the fibrillar centre. Nucleolar area was quantified using Image J and fibrillarin staining for at least 250 cells per experiment. Graph depicts the mean (±s.e.m) of three experiments. **(I)** rDNA transcription and nucleolar size were monitored over time in SW480 cells using qRT-PCR for the 47S transcript (as above) and ImageJ analysis of area devoid of DAPI staining (as a marker for nucleoli). At least 200 cells from 10 fields of view were analysed for nucleolar area. Graph depicts the mean of three experiments (±s.e.m). *** P<0.001. Actin and Cu/ZnSOD act as loading controls throughout. Scale bars=10μM. *P* values throughout are compared to the respective control and were derived using a two tailed students Ttest. See Supplemental Figure 2 for additional cell lines and supporting data.

TIF-IA qRT-PCR and cyclohexamide run on assays indicated the reduction in TIF-IA protein in response to aspirin was not a consequence of reduced gene transcription, but caused by increased protein turnover (Figures 2*E* and *F* and Supplemental figure 2*C*). Exogenously expressed TIF-IA was also rapidly degraded in response to aspirin (Supplemental Figure 2*D*). In contrast, the agent had no effect on levels of UBF or POLR1A (Figure 2*A*).

Classic hallmarks of nucleolar stress are inhibition of rDNA transcription, segregation of nucleolar marker proteins and reduced nucleolar area (1). Given the role of TIF-IA in maintaining nucleolar structure(41), we investigated these hallmarks. We found aspirin-mediated TIF-IA degradation was associated with a significant decrease in rDNA transcription, as indicated by 47S qRT-PCR and 5-fluorouridine (FUrd) run on assays (Figures 2*G* and Supplemental figure 2*E*). It was also associated with nucleolar segregation, as evidenced by relocation of all three components of the tri-partite nucleolar sub-structure to the periphery of the organelle (Figure 2*H* and Supplemental figure 2*F*). However, in contrast to actinomycinD which as expected, caused a significant *reduction* in nucleolar area, aspirin induced a significant *increase* in nucleolar area (Figure 2*H*). This increase was an early response to the agent, paralleling degradation of TIF-IA and inhibition of rDNA transcription (Figures 2*C* and *I*).

### Generality of TIF-IA degradation in response to stress stimuli of the NF-κB pathway

We next examined the generality these effects with regards to stress stimuli of NF-κB and found that like aspirin, UV-C exposure induced a rapid depletion of TIF-IA (Figures 3*A* to *C*). Although this depletion was transient in SW480 cells, it still preceded degradation of IκB and was paralleled by enlargement of nucleoli (Figures 3*A* to *C*). In contrast to UV-C, the DNA damaging agent camptothecin, which has previously been shown to induce nucleolar stress (Table S1), actually increased cellular levels of TIF-IA (Figure 3*D*). It also had a minimal effect on nucleolar structure (Figure 3*D*). Similarly, actinomycinD, BMH-21, CX5461 and TNF did not significantly alter TIF-IA levels (Figure 3*E*).

**Figure 3.**
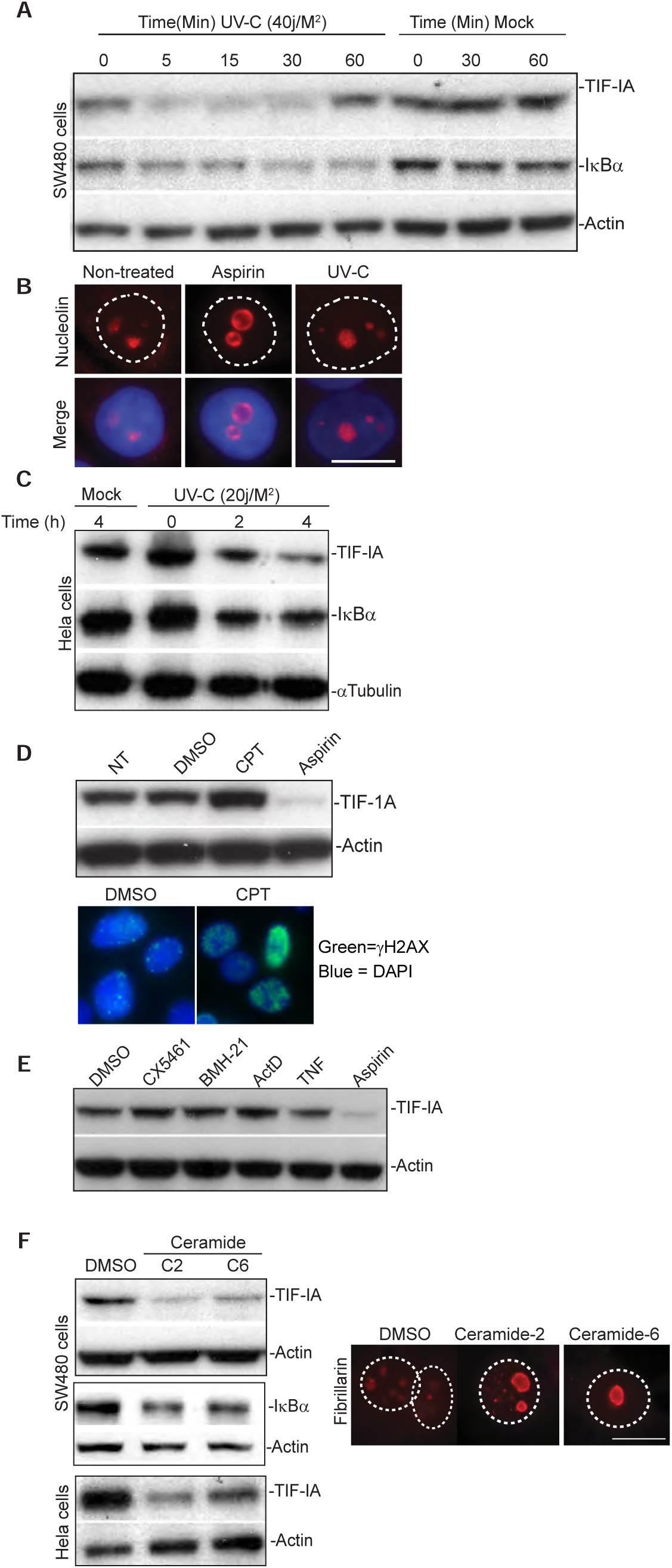
TIF-IA degradation in response to multiple stress stimuli of NF-κB (A and C) SW480 or Hela cells were mock or UV-C (20 or 40 J/m^2^) irradiated. Following the times specified, immunoblots was performed on WCL with the indicated antibodies. **(B)** Immunomicrograph (x63) showing increased nucleolar area (as depicted by nucleolin staining) in SW480 cells in response to aspirin (3mM, 16h) or UV-C (40j/m^2^, 2h). **(D)** SW480 cells were treated with carrier, 10μM camptothecin (CPT) or aspirin (3mM) for 16h. Top: Western blot was performed with the indicated antibodies on WCL. Bottom: γH2AX immunocytochemistry confirmed DNA damage in response to CPT. **(E)** SW480 cells were treated with DMSO (carrier), CX5461 (500nM), BMH-21 (4μM), ActinomycinD (ActD, 50ng/ml) or aspirin (10mM) for 4 hours, or TNF (10ng/ml) for 30mins, WCL were examined by western blot using the antibodies indicated. (**F**) SW480 or Hela cells were treated with carrier (DMSO), ceramide-2 (C2, 10uM)) or ceramide-6 (C6, 10uM) for 16h. WCL were analysed by western blot with the indicated antibodies. Representative immunomicrographs (X63) show the localisation of fibrillarin in SW480 cells. Scale bars=10μM. Actin or α tubulin act as loading controls.

Aspirin and UV-C generate ceramide, a crucial lipid second messenger that is a potent stimulus of the NF-κB pathway(42-44). When we tested the C2 and C6 soluble forms of ceramide, which were used to mimic the lipid increase observed in response to these and other stresses, we found that they also induced degradation of TIF-IA (Figures 3*F* and *G*). Furthermore, this TIF-IA degradation occurred in association with increased nucleolar area and degradation of IκB (Figures 3*F*).

Together, these data suggest that specific stresses, through the generation of ceramide, target TIF-IA for degradation, and that this degradation may cause distinctive changes in nucleolar structure and activation of NF-κB. To explore this possibility, we set out to elucidate the mechanism underlying stress-mediated TIF-IA degradation, again using aspirin as a model stimulus.

### Proteasomal/lysosomal degradation of TIF-IA

Nguyen et al previously demonstrated that basal TIF-IA turnover is regulated by proteasomal degradation, facilitated by the E3 ligase, MDM2(20). Therefore, we firstly investigated this pathway. We found the proteasome inhibitor, MG132, and the MDM2 inhibitor, nutlin-3, blocked basal degradation of TIF-IA as previously reported (20) (Figure 4*A*). However, neither inhibitor blocked aspirin-mediated degradation (Figure 4*A*). Indeed, this was enhanced in the presence of Nutlin-3 (Figure 4*A*). Similar results were obtained with the highly specific proteasome inhibitor, lactacystin (Supplemental figure 3*A*).

**Figure 4.**
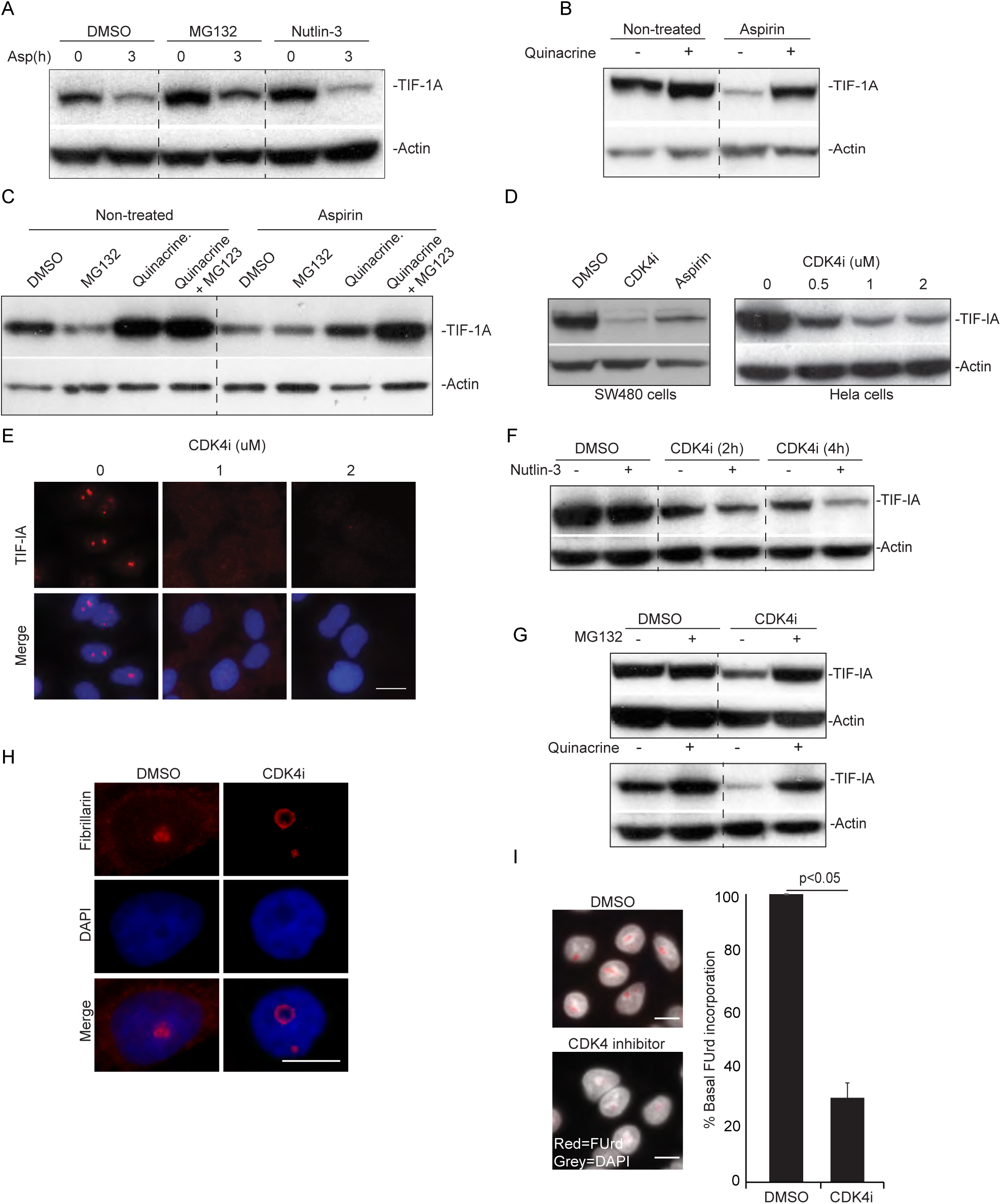
CDK4 inhibition mimics stress effects on TIF-IA and nucleoli. **(A to C)** Aspirin-mediated TIF-IA degradation is independent of MDM2 and dependent on lysosomal and proteasomal pathways. SW480 cells were pre-treated for 2h with DMSO (carrier), the MDM2 inhibitor, Nutlin3 (5μM), MG132 (25μM), Quinacrine (25μM) or MG132 (10μM) plus quinacrine. Following aspirin treatment (10mM, 4h or as indicated), TIF-IA levels were monitored by Western blot analysis. **(D to F)** CDK4 inhibition induces degradation of TIF-IA. **(D and E)** SW480 or Hela cells were treated with DMSO (carrier), aspirin (3mM, 16h) or the small molecule CDK4 inhibitor, 2-bromo-12,13-dihy-dro-indolo[2,3-a]pyrrolo[3,4-c] carbazole-5,7(6H)-dione (CDK4i, 2uM or as indicated). **(D)** Anti-TIF-IA immunoblot performed on WCL. **(E)** Immunomicrographs (63X) demonstrating the levels and localisation of TIF-IA. **(F and G)** SW480 cells were pre-treated with Nutlin-3, MG132 or Quinacrine as above, prior to exposure to CDK4i (2μM) for 2h, or times specified. Western blot analysis was performed on WCL with the indicated antibodies. **(H and I)** Modulation of nucleolar structure and function by CDK4 inhibition. SW480 cells were treated with carrier (DMSO) or CDK4i (2μM) for 16h. **(H)** Immunomicrograph (63X) demonstrating re-localisation of fibrillarin in response to CDK4i. **(I)** Left: Immunomicrographs (40X) depicting cells subjected to fluouridine (FUrd) run on assays. Right: Images were captured and analysed for FUrd incorporation using ScanR image analysis software. The results are presented as the percentage incorporation compared to control. The mean (± s.e.m) of at least 1000 cells in three independent experiments is shown. Scale bar =10μm. Actin acts as a loading control throughout. *P* values were derived using a two tailed students Ttest. See also Supporting Figure 3.

Next, we investigated lysosomal degradation and found that inhibiting lysosome activity with quinacrine caused a substantial increase in basal TIF-IA (Figure 4*B*). Quinacrine also partially reduced aspirin-mediated TIF-IA degradation, implicating lysosomal involvement (Figure 4*B*). Given the redundancy in degradative pathways, we inhibited the lysosome and proteasome together. At the lower concentrations used in combination studies, MG132 alone caused a reduction in TIF-IA. However, this was completely blocked in the presence of quinacrine (Figure 4*C* lane 4). Aspirin-mediated TIF-IA degradation was also further abrogated by the inhibitor combination (Figure 4*C*, lane 8). Based on these data, we concluded that TIF-IA is degraded in response to stress in a manner dependent on both the proteasome and lysosome and that the upstream pathway to this degradation differs from basal turnover.

### CDK4 inhibition mimics stress effects on TIF-IA and nucleoli

A common early response to aspirin, UV-C and ceramide is reduced activity of CDK4/6 (45). Since CDK4 targets components of the PolI complex, we considered that inhibition of this kinase may lie upstream of stress effects on TIF-IA. To test this possibility, we utilised a highly specific, small molecule inhibitor of CDK4 (CDK4i) which we have previously shown stimulates the NF-κB pathway(45).

Immunocytochemistry and immunoblot analysis demonstrated that CDK4i induces a substantial, dose dependent reduction in TIF-IA ( Figures 4*D* and *E*). This effect was evident in multiple cell types (Figure 4*D*). It was also observed in response to an independent CDK4 inhibitor, PD0332991, indicating specificity (Supplementary figure 3*B*). Like aspirin, CDK4i-mediated TIF-IA degradtion was enhanced in the presence of nutlin-3 (Figure 4*F*). It was also blocked by proteasome and lysosomal inhibitors, again suggesting dependency on both degradative mechanisms (Figures 4*G*). Furthermore, it was paralleled by the same distinct nucleolar phenotype as stress stimuli of the NF-κB pathway i.e increased nucleolar size, segregation of nucleolar marker proteins and inhibition of rDNA transcription (Figures 4*H* and *I*).

### Identification of a role for UBF S484A and p14ARF in TIF-IA degradation

CDK4 regulates rRNA transcription by phosphorylating UBF (46) and so, we considered UBF may be involved in this TIF-IA degradative mechanism. Indeed, siRNA silencing of UBF significantly abrogated CDK4i-mediated TIF-IA degradation (Figures 5*A* and *B*). Silencing of UBF also abrogated TIF-IA degradation in response to aspirin, although the effect was not as pronounced with this less specific agent (Figures 5*A* and *B*). In contrast to UBF, silencing POLR1A did not perturb TIF-IA degradation in response to either agent, suggesting specificity (Figure 5*A*).

**Figure 5.**
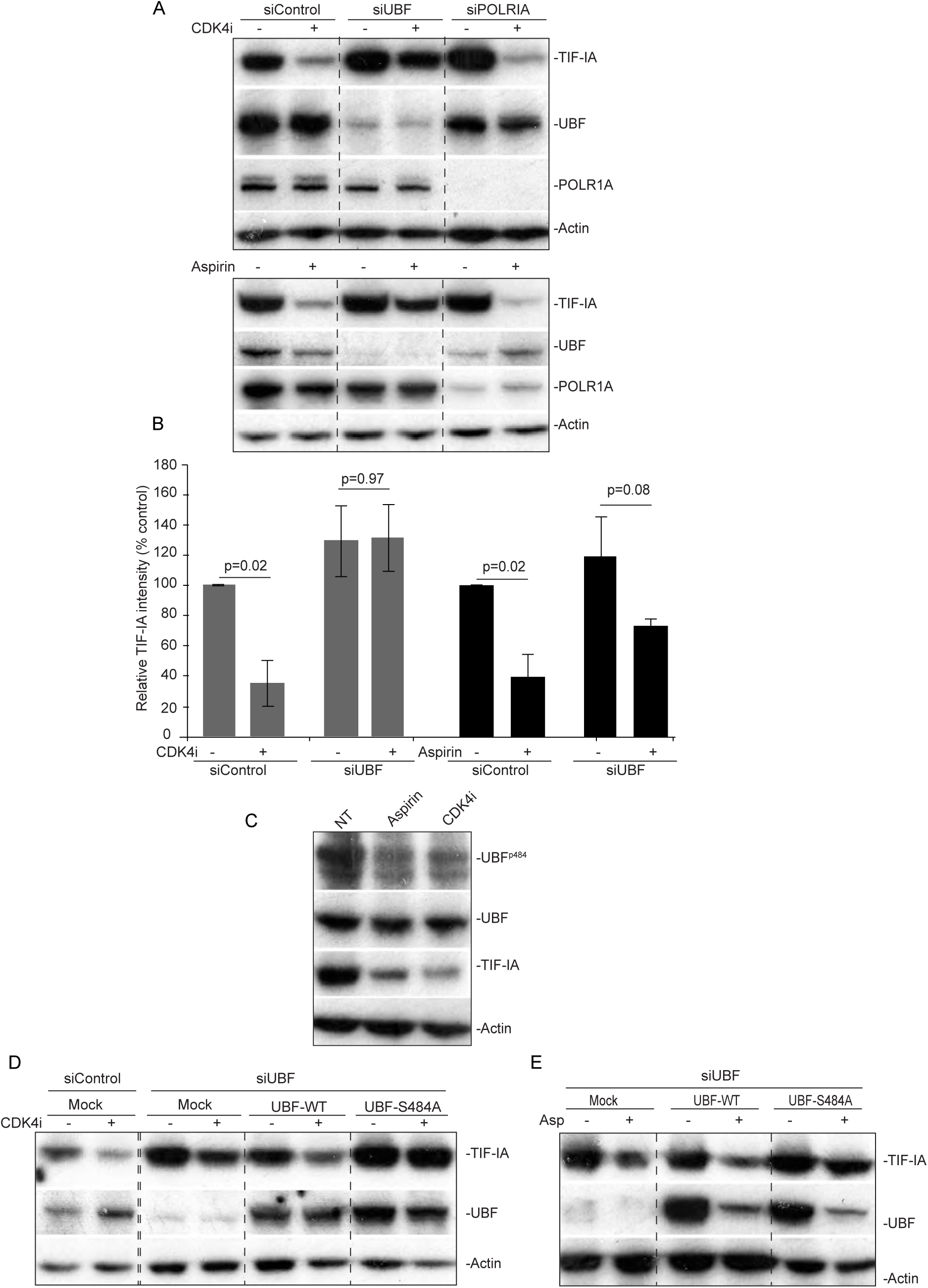
Identification of a role for UBF S484 in TIF-IA degradation. **(A and B)** UBF is required for CDK4i-and aspirin-mediated TIF-IA degradation. SW480 cells were transfected with control, UBF or POLRIA siRNA. Seventy two hours later cells were treated (+) with CDK4i (2uM, 4h), aspirin (3mM 16h) or the equivalent carriers (-). **(A)** Western blot analysis was performed with the indicated antibodies. **(B)** TIF-IA intensity (relative to actin) was determined for each condition using ImageJ analysis. Results are presented as the percentage relative TIF-IA compared to carrier treated, siControl. Mean (± s.e.m) is shown for 2 (CDK4i) and 3 (aspirin) experiments. **(C to E)** Identification of a role for residue 484 of UBF. **(C)** SW480 cells were treated with aspirin and CDK4i as above. Western blot analysis was performed with an antibodies to UBF phosphorylated at S484 and native UBF. **(D and E)** SW480 cells were transfected with control or UBF siRNA then either mock transfected or transfected with Flag-UBF-wild type (WT) or a phospho-mutant, flag-UBFS484A. Eight hours later, transfected cells were treated with CDK4i (2uM, 16h) or aspirin (3mM, 16h). Immunoblot was performed with the indicated antibodies.

CDK4 phosphorylates UBF on Serine 484 so we next explored the role of this residue(46). Figure 5C demonstrates that both aspirin and CDK4i cause a decrease in UBF S484 phosphorylation. To determine whether this decrease is essential for TIF-IA degradation we utilised a mutant that cannot be phosphorylated at this site (Flag-UBF S484A). SW480 cells were depleted for UBF then transfected with plasmids expressing either wild type protein (Flag-UBF-WT) or the Flag-UBF-S484A mutant. Again, we found that silencing of UBF abrogated CDK4i-mediated degradation of TIF-IA (Figure 5*D*). Expression of wild type UBF rescued this effect, confirming an important role for this protein. If de-phosphorylation of UBF at S484 was important for TIF-IA degradation, we would expect that the phospho-mutant would mimic the effects of CDK4i on TIF-IA, or at least enhance CDK4i-mediated degradation of the protein. However, contrary to this expectation, expression of flag-UBF-S484A actually blocked TIF-IA degradation in response to CDK4i and aspirin (Figures 5*D* and *E*). Together, these data indicate a crucial role for UBF, and in particular residue 484, in stress/CDK4i-mediated degradation of TIF-IA. However, the phosphorylation status of this residue does not appear to be critical.

p14ARF is a nucleolar tumour suppressor that regulates rRNA synthesis by targeting S484 of UBF (47,48). It is also known to influence NF-κB signalling (49) and therefore, we considered it may play a role. Indeed, immunoprecipitation assays revealed that TIF-IA complexes with P14ARF in response to aspirin in a time and dose dependent manner that parallels degradation of the protein (Figures 6A and Supplementary 4A). Furthermore, immunoblot analysis indicated that siRNA silencing of p14ARF blocks CDK4i and aspirin-mediated degradation of TIF-IA, while over-expression enhances this effect (Figures 6B to D). Aspirin alone did appear to cause a dose dependent reduction in p14ARF levels (Figure 6B). However, detailed time course studies revealed that TIF-IA degradation precedes loss of p14ARF (Supplementary figure 4*B*). Immunocytochemical analysis confirmed that aspirin and CDK4i cause a significant reduction in nuclear TIF-IA in cells transfected with control siRNA, but not in cells transfected with p14ARF siRNA (Figures 6E and F). Interestingly, in the absence of p14ARF, TIF-IA remained within nucleoli following aspirin and CDKi exposure suggesting this protein may play a role in nucleolar export (Figures 6E and F).

**Figure 6.**
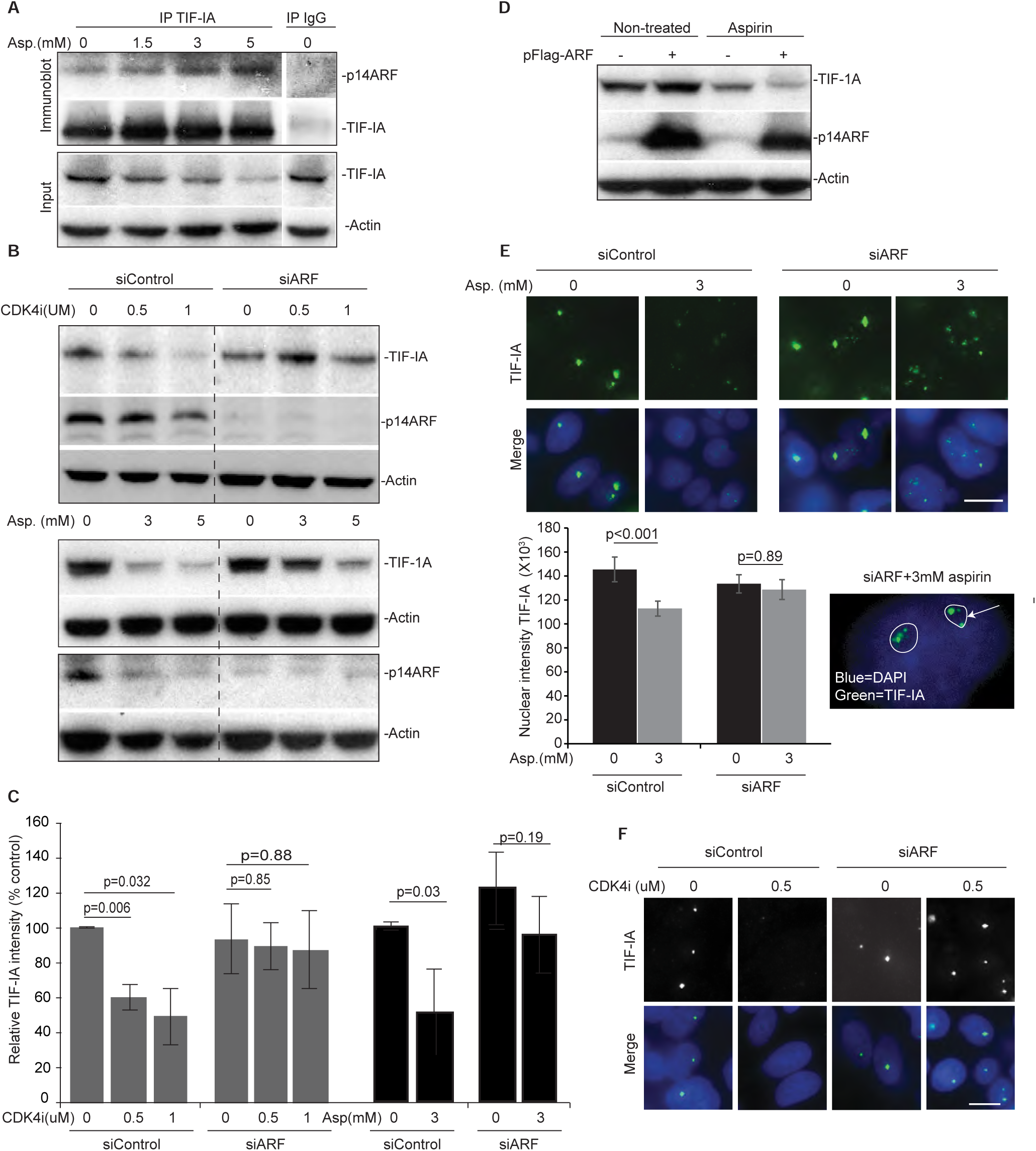
Identification of a role for p14ARF in TIF-IA degradation. **(A)** TIF-IA interacts with p14ARF in response to aspirin. SW480 cells were treated with 0-5mM aspirin for 16h. Immunoprecipitation was carried out on WCL using antibodies to TIF-IA and IgG control. Precipitated proteins were subjected to western blot analysis with the indicated antibodies. Input levels are shown. **(B to F)** Silencing of P14ARF is required for stress-mediated degradation of TIF-IA. (**B, C, E and F**) SW480 cells were transfected with control or p14ARF siRNA (siARF) then treated for 16h with CDK4i (0-1μM) or aspirin (0-5mM). **(B)** Immunoblot analysis was performed on WCL with the indicated antibodies. **(C)** TIF-IA levels relative to actin were quantified using ImageJ analysis. Graph shows the mean (± s.e.m) compared to non-treated, siControl. N=3. **(E and F)** Immunomicrographs (X63) show the levels and localisation of TIF-IA in fixed cells. **(E)** IPlab software quantified nuclear (as depicted by DAPI staining) intensity of TIF-IA. Data are the mean (±s.e.m) of >150 nuclei. N=3. Inset shows nucleolar (outlined) TIF-A in p14ARF transfected cells treated with aspirin. **(D)** SW480 cells were transfected with pcDNA3 control plasmid (-) or pcDNA3-p14ARF (+) then either non-treated or treated with aspirin (3mM 16h). Immmunoblot was performed on WCL with the indicated antibodies. Actin acts as a loading control throughout. Scale bar =10μm. P values were derived using a 2 tailed students Ttest. See also supporting Figure 4.

### Inhibiting TIF-IA degradation inhibits activation of the NF-κB pathway

Using p14ARF siRNA as a tool to block TIF-IA degradation, we next examined the role of this degradation in stress effects on NF-κB signalling and nucleolar structure. Figures 7*A* to C demonstrate that blocking TIF-IA degradation in this manner significantly inhibits aspirin-mediated inhibition of rDNA transcription and enlargement of nucleoli. Furthermore, the significant degradation of IκB and nuclear/nucleolar accumulation of RelA observed in response to aspirin in control siRNA transfected cells was blocked in cells transfected with siRNA to p14ARF (Figures 7B, *D* and *E*). Aspirin-mediated apoptosis was also abolished by silencing of p14ARF, which would be expected given our data showing NF-κB pathway activation is required for the apoptotic effects of the agent(31) (Figure 7*F*). Similar results were obtained with CDK4i in that p14ARF silencing abrogated CDK4i-mediated nucleolar enlargement, degradation of IκB and nuclear accumulation of RelA (Figures 8*A* and *B*).

**Figure 7.**
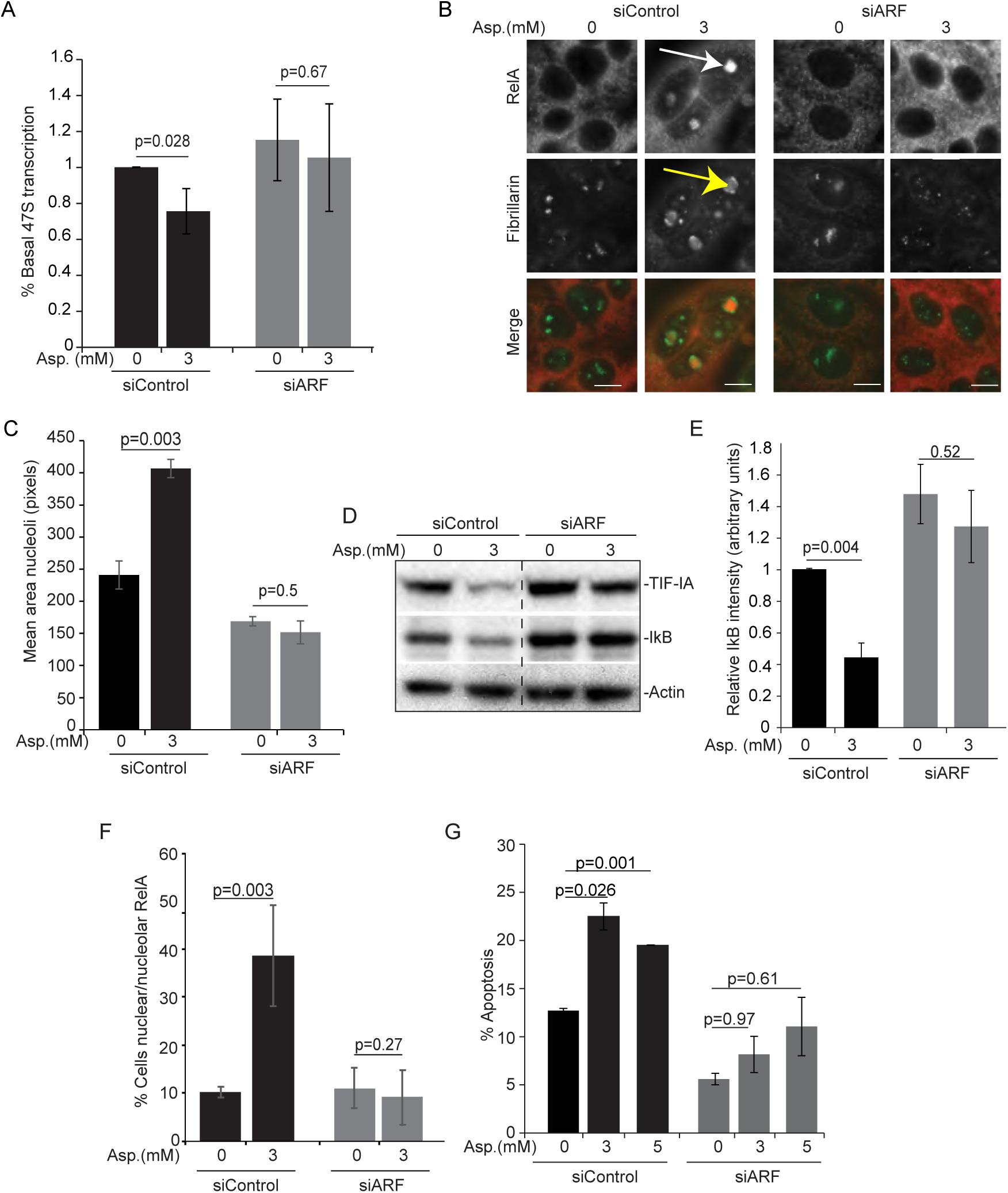
Blocking TIF-IA degradation inhibits aspirin effects on nucleoli and the NF-κB pathway. **(A to G)** Blocking TIF-IA degradation, using siRNA to p14ARF, abrogates aspirin-mediated inhibition of rDNA transcription, nucleolar enlargement, degradation of IκB, nuclear/nucleolar translocation of RelA and apoptosis. SW480 cells were transfected with control or p14ARF siRNA as in Figure 6 then treated with aspirin (Asp.) at the concentrations specified. **(A)** qRT-PCR with primers for the 47S pre-rRNA transcript measured levels of rRNA transcription. GAPDH was used to normalise. Results are presented as the percentage of relative 47S transcription compared to the equivalent 0mM control for each siRNA. Mean (± s.e.m) of 3 is shown. **(B)** Immunocytochemistry was performed on fixed cells with the indicated antibodies. Arrows indicate nucleolar RelA (white) and enlarged, segregated nucleoli (yellow). **(C)** Nucleolar area was quantified in at least 150 cells using IPlab software with fibrillarin as a nucleolar marker. Mean (± s.e.m) is shown. N=3. **(D)** Immunoblots demonstrating cytoplasmic levels of IκBα. **(E)** ImageJ software measured IκBα intensity relative to actin. The results are the mean of 3 experiments ±s.e.m. **(F)** Immunocytochemistry was performed as in B. The percentage of cells in the population showing nucleolar RelA was quantified. At least 6 fields of view and >100 cells were analysed per experiment. The mean ± s.e.m is shown. N=3. **(G)** Annexin V apoptosis assays were performed. The percentage of cells undergoing apoptosis was determined by fluorescent microscopy. At least 200 cells were analysed for each sample. The results are the means of two independent experiments ± s.e.m. Actin acts as a loading control. Scale bar =10μm. P values were derived using a 2 tailed students Ttest.

**Figure 8.**
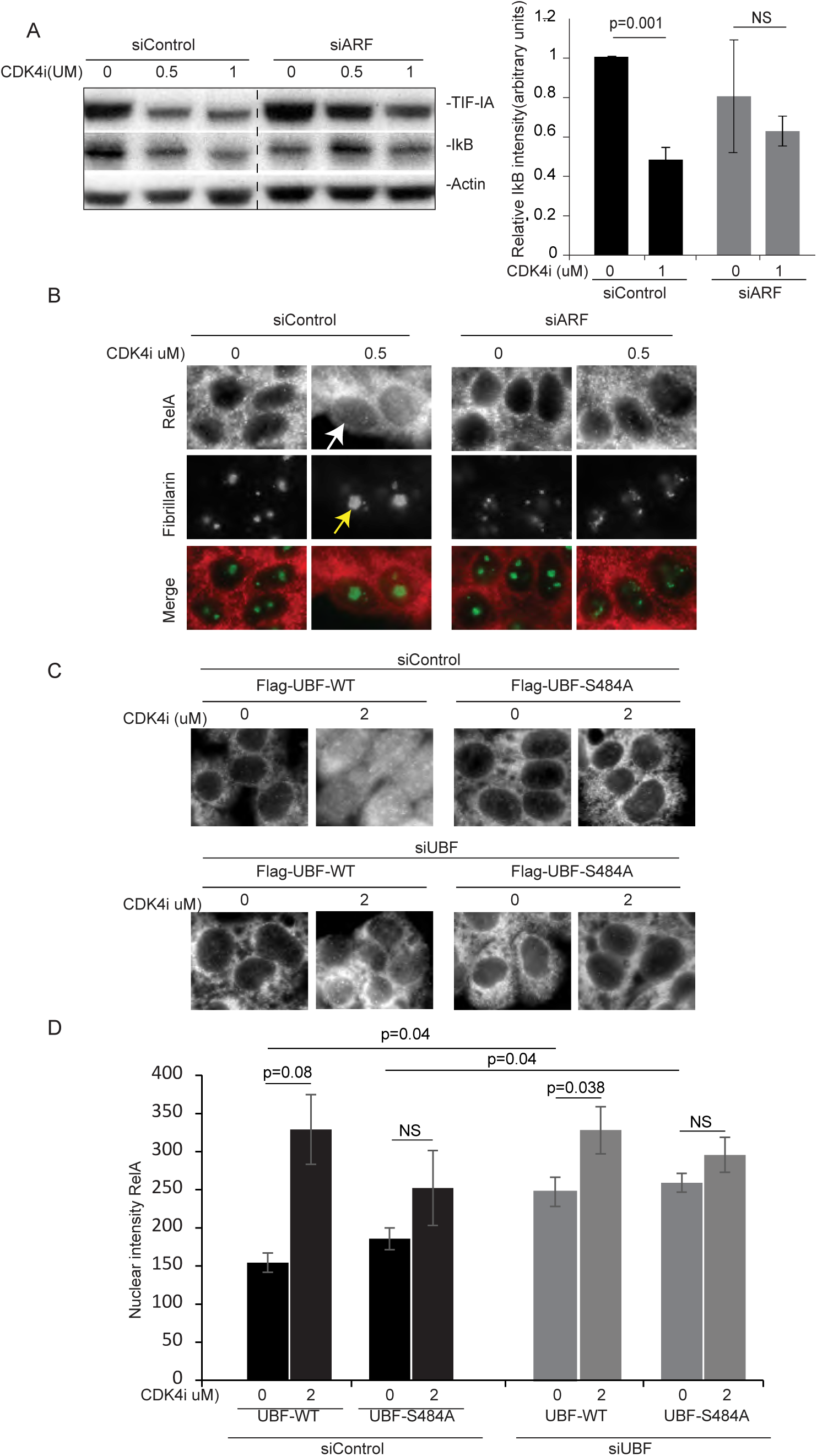
Blocking TIF-IA degradation inhibits CDK4i effects on nucleoli and the NF-κB pathway. **(A and B)** SW480 cells were transfected with control or p14ARF siRNA as above then treated with CDK4i at the concentrations specified. **(A)** Immunoblot demonstrating cytoplasmic levels of IκBα. Right: ImageJ software measured IκBα intensity relative to actin. The results are the mean of 2 experiments ±s.e.m. **(B)** Immunocytochemistry was performed on fixed cells with the indicated antibodies. Arrows indicate increased nuclear RelA (white) and nucleolar area (yellow) in response to CDK4i in siControl transfected cells. **(C)** SW480 cells were transfected with the indicated siRNA species alongside Flag-UBF WT or S484A (as in Figure 5D). Following CDK4i treatment (0-2μM), immunocytochemistry was performed on fixed cells with antibodies to RelA. ImageJ quantified the nuclear (determined by DAPI) intensity of RelA. Graph depicts the mean of at least 200 cells/experiment (± s.e.m). Actin acts as a loading control throughout. Scale bar =10μm. P values were derived using a 2 tailed students Ttest.

Next, we utilised the UBF-S484A mutant to investigate the link between TIF-IA degradation and NF-κB signalling. SW480 cells were transfected with control or UBF siRNA prior to overexpression of Flag-UBF-WT or -S484A. Quantitative immunocytochemistry demonstrated that expression of the mutant alone caused a significant increase in nuclear RelA (Figure 8C), in keeping with our data showing disruption of the PolI complex activates the NF-κB pathway (Figure 1A). Quantitative immunocytochemistry also demonstrated that the significant increase in nuclear RelA observed in response to CDK4i in cells expressing WT-UBF was severely abrogated in those expressing UBF-S484A (Figures 8C and D).

Together, these data reveal a new pathway by which stresses act on the PolI complex that involves CDK4-UBF(S484)/p14ARF facilitated degradation of TIF-IA. They also identify atypical changes in nucleolar architecture and activation of the NF-κB pathway as a novel downstream consequence of this specific PolI complex disruption (Figure 9A).

**Figure 9.**
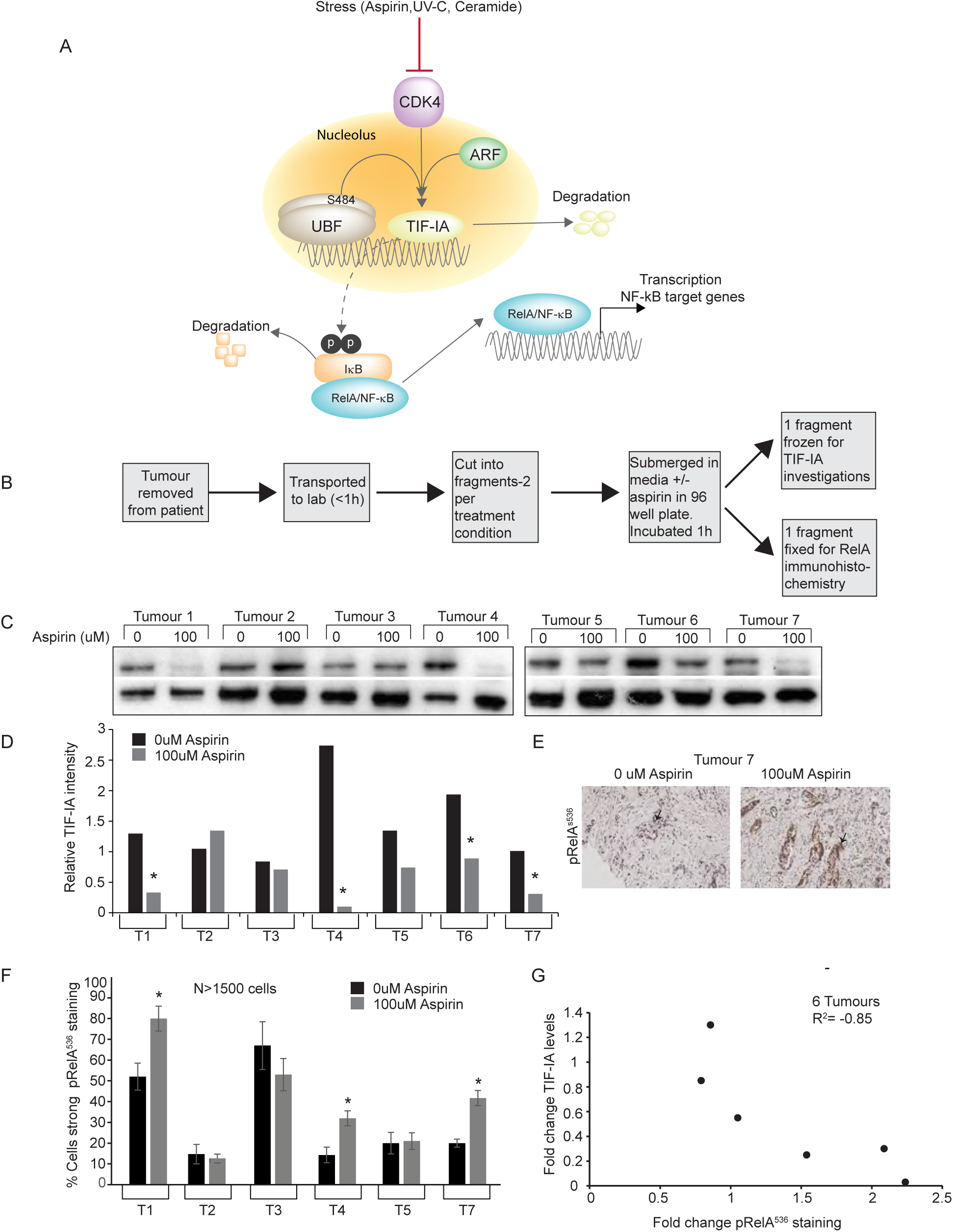
TIF-IA degradation correlates with NF-κB pathway activation in human clinical samples. **(A)** Proposed model. Inhibition of CDK4 kinase activity by specific stresses induces degradation of TIF-IA in a UBF^S484^/p14ARF dependent manner. The consequent disruption of the PolI pre-initiation complex causes a protein or pathway to be triggered (as yet unknown) which stimulates phosphorylation and degradation of IκBα, nuclear translocation of RelA and ultimately, transcription of gene programme that alters cell phenotype. We propose the nature of this transcriptional program is cell type and stimulus dependent. **(B)** Diagram depicting workflow of *ex vivo* culture. Resected colorectal tumour biopsies were immediately transferred to the lab, washed, immersed in culturing media in 96 well plates then exposed to 0 or 100uM aspirin for 1h. One piece of tissue was fixed for immunohistochemistry while another was frozen for protein analysis. This was carried out for 7 patients. **(C)** Immunoblot was performed on WCL with the indicated antibodies. **(D)** ImageJ quantified TIF-IA intensity relative to actin. *= Tumours showing a greater than 2 fold decrease in relative levels of TIF-IA in response to aspirin. **(E and F)** Immunohistochemistry was performed on sections from paraffin embedded tissue with antibodies to RelA^p536^. **(E)** An example immunomicrograph. Arrows indicate epithelial cells. **(F)** Images were digitised then Leica QWin plus image analysis software used to quantify the nuclear RelA^p536^ intensity. Three distinct areas of tissue and at least 1500 cells were analysed per section. Data presented are the % cells showing moderate+strong RelA^p536^ staining, as indicated by image analysis software. * = Significant (P<0.05) difference between the % stained cells in treated and non-treated sections. (**G)** Graph showing the relationship between aspirin-induced changes in TIF-IA and RelA^p536^ staining for 6 individual tumours.

### Relationship between TIF-IA degradation and stimulation of NF-κB signalling in human clinical samples

Overwhelming evidence indicates that aspirin has anti-tumour activity and the potential to prevent colorectal and other cancers (50,51). To investigate the clinical significance of our results with regards to this activity, and to determine whether there is a link between PolI complex disruption and NF-κB pathway activation in a whole tissue setting, we treated biopsies of fresh, surgically resected human colorectal tumours with pharmacological doses (0-100 μM, 1h) of aspirin *ex vivo* (Figure 9A). This aspirin concentration is comparable to salicylate levels we measured in plasma from patients given a short course of analgesic doses of aspirin (31). It is also well within the reported therapeutic range (0.1-3mM).

Western blot analysis revealed low dose aspirin induces TIF-IA degradation (as defined by a > 2 fold reduction in protein levels) in 4/7 (57%) tumours exposed *ex vivo* to the agent (Figure. 9*B*). Furthermore, quantitative immunohistochemistry with antibodies to pRelA^536^ indicated *ex vivo* exposure to low dose aspirin induced NF-κB pathway activation in 3/6 tumours (Figure 7*C*). Importantly, for individual tumours, there was a very strong inverse correlation (r2=-0.85, n=6) between aspirin effects on TIF-IA and RelA^536^ phosphorylation (Figure 9*D*). That is, the greater the loss of TIF-IA, the greater the increase in NF-κB pathway activation. These data confirm that aspirin causes PolI complex disruption and activates the NF-κB pathway in primary human tumours and suggests a strong relationship between these two events in a whole tissue setting. These data have far reaching implications for understanding of the anti-tumour effects of this agent.

## Discussion

The work presented here has great significance as we identify a novel pathway by which PolI activity is inhibited by stress and for the first time, identify NF-κB activation as a downstream consequence of nucleolar stress response (Figure 9A). We demonstrate the relevance of this pathway *in vivo* using human clinical samples and show it may contribute to the anti-tumour effects of aspirin. These data shed new light on the mechanisms by which nucleoli sense stress and coordinate the downstream consequences.

### Identification of an NF-κB nucleolar stress response pathway

The paradigm of nucleolar stress response is inhibition of rDNA transcription leading to stabilisation of p53 (8). Here we demonstrate that specifically disrupting the PolI complex activates the cytoplasmic NF-κB pathway, but that this effect is not mimicked by potent inhibitors of rDNA transcription (Figure 1). We also demonstrate that certain NF-κB stimuli induce degradation of the PolI complex component, TIF-IA (Figures 2-4). Using two independent approaches, we demonstrate that blocking this degradation blocks the effect of these agents on the NF-κB pathway (Figures 7 and 8). Finally, we demonstrate a strong correlation between loss of TIF-IA and activation of NF-κB in a whole tissue setting (Figure 9). Together, these data provide powerful evidence for an independent nucleolar stress response pathway that is characterised by degradation of TIF-IA leading to activation of NF-κB. In support of this pathway being distinct from classical nucleolar stress, stimulation is associated a distinct nucleolar phenotype. That is, inhibition of rDNA transcription alongside *increased* nucleolar area and segregation of nucleolar marker proteins.

We are currently exploring the mechanism that links TIF-IA degradation to stimulation of the NF-κB pathway. We are working on the premise that it differs from the mechanism leading to p53 stabilisation (cytoplasmic release of RPL11/RPL5), as the p53 nucleolar stress response pathway is triggered by CX5461 and BMH21 (36,52), which had no effect on NF-κB-driven transcription. CK2 is an excellent candidate protein as it is found as part of the PolI complex (23) and phosphorylates IκBα in response to UV-C(24). Another kinase of interest is NIK (NF-κB inducing kinase), which acts upstream of the IkappaB kinase (IKK) complex and is known to shuttle through nucleoli (53). The ribosomal proteins L3 and S3 have also been shown to complex with IκB and modulate NF-κB activity (29,53,54).

Depletion of PolI components, CDK4 inhibition and stress stimuli of NF-κB all induce cell cycle arrest(55,56). Therefore, we did consider that it may be this that stimulates NF-κB signalling, rather than loss of PolI complex integrity *per se*. However, we have previously shown that inhibiting each stage of the cell cycle does not mimic the effects of CDK4i on nucleoli or on the NF-κB pathway(56). Furthermore, we found that p14ARF siRNA blocks aspirin-mediated activation of the NF-κB pathway, but not degradation of cyclinD1 (data not shown). Therefore, we conclude that in response to specific stimuli, activation of NF-κB is a consequence of TIF-IA degradation.

Given that NF-κB stress stimuli inhibit rRNA transcription, it is highly likely that they also stabilise p53. We have previously demonstrated that stimulation of the NF-κB pathway is absolutely necessary for aspirin-mediated apoptosis, but that p53 is dispensable (31,57). Here we show that blocking TIF-IA degradation blocks aspirin-mediated stimulation of the NF-κB pathway and apoptosis. Therefore, we believe that in this context, the phenotypic response to nucleolar stress is governed by the NF-κB pathway.

Our data indicate TIF-IA degradation is associated with reduced rRNA transcription, but increased nucleolar area. This phenotype would appear to contradict the belief, which is used widely by pathologists, that increased nucleolar size is a marker for enhanced rRNA transcription. In keeping with our findings, Fatyol et al found that MG132 induces a significant increase in nucleolar volume while inhibiting rRNA transcription and mediating cell death(58). Similarly, Bailly et al found that the NEDD8 inhibitor, MLN4924, causes an increase in nucleolar size while inducing cell death, although in this case there was no effect on rRNA transcription(59). These data bring into question the use of nucleolar size as an absolute marker of cell proliferation in diagnostic pathology.

### Identification of a novel pathway to TIF-IA inactivation

It is well documented that the phosphorylation status of TIF-IA is modulated in response to environmental and cytotoxic stress so that rates of rRNA transcription can be adjusted accordingly (37). Here we identify an alternative mechanism by which stress can act on the PolI complex involving degradation of TIF-IA. We also present several lines of evidence to suggest that this degradation is dependent upon CDK4 inhibition and UBF/p14ARF. Firstly, we demonstrate that stress effects on TIF-IA can be mimicked by two highly specific inhibitors of CDK4. We also demonstrate siRNA depletion of UBF or p14ARF blocks aspirin and CDK4 i-mediated degradation of TIF-IA, while overexpression of either of these proteins enhances this effect. Finally, we show that expression of UBF S484A, a mutant that cannot be phosphorylated at this site, blocks aspirin and CDK4i-mediated TIF-IA degradation. How CDK4 inhibition, UBFS484 and p14ARF combine to promote the degradation of TIF-IA is complex and out-with the scope of this study. Given that the UBFS484 phospho-mutant blocked TIF-IA degradation, we suggest that inhibition of CDK4 kinase activity is not directly affecting this site, but that another CDK4 target is involved. Retinoblastoma protein, which binds to UBF in a hypophosphorylated state, is a good candidate and the subject of ongoing investigation (60).

### Aspirin degradation of TIF-IA:Identification of a novel mechanism of action

Despite the burden of evidence indicating aspirin use can prevent colorectal cancer, the agent cannot be recommended for this purpose due to its significant side effect profile. Hence, efforts are now focussed on understanding the mechanism by which aspirin acts against colorectal cancer cells to identify markers of response and targets for safer, more effective alternatives. Uncontrolled rDNA transcription is a hallmark of cancer and contributes to tumour growth by allowing de-regulated protein synthesis and uncontrolled activity of nucleolar cell growth/death pathways(36,61). Hence, inhibition of the PolI complex is emerging as an attractive therapeutic strategy. Here we show for the first time that aspirin induces degradation of TIF-IA and inhibits rDNA transcription. This is an extremely exciting, novel mechanism of action that could have a significant impact in defining patients that would benefit from aspirin therapy.

In summary, the data presented here open up new avenues of research into nucleolar regulation of NF-κB signalling and regulation of TIF-IA stability under stress. They also shed further light on the complex mode of action of aspirin and related non-steroidal anti-inflammatory drugs (NSAIDs).

## Funding

The work was supported by grants from the WWCR [formally AICR grant number:10-0158 to LS], Rosetrees Trust [Grant numbers A631, JS16/M225 to LS], BBSRC [Grant number BB/H530362/1 to LS for SN] and MRC [Grant number MR/J001481/1]. JC and IL were supported by University of Edinburgh scholarships.

## Acknowledgements

We would like to thank RT. Hay (University of Dundee), A Lamond (University of Dundee), I Grummt (German Cancer Centre), H Bierhoff (German Cancer Centre), M Laiho (Johns Hopkins University School of Medicine) and B McStay (NUI Galway) for providing tools and reagents. We would also like to thank Nick Hastie, Arkadiusz Welman and Wendy Bickmore for critically reading the manuscript. C. Nicol provided help with figure preparation, ECMC Edinburgh with tissue collection and M. Walker with general laboratory support.

## References

1. Boulon, S., Westman, B.J., Hutten, S., Boisvert, F.M. and Lamond, A.I. (2010) The Nucleolus under Stress. Mol. Cell, 40, 216–227.

2. Grummt, I. (2013) The nucleolus-guardian of cellular homeostasis and genome integrity. Chromosoma.

3. Boisvert, F.M., van Koningsbruggen, S., Navascues, J. and Lamond, A.I. (2007) The multifunctional nucleolus. Nat. Rev. Mol. Cell Biol, 8, 574–585.

4. Mayer, C. and Grummt, I. (2005) Cellular stress and nucleolar function. Cell Cycle, 4, 1036–1038.

5. Boisvert, F.M., Lam, Y.W., Lamont, D. and Lamond, A.I. (2010) A quantitative proteomics analysis of subcellular proteome localization and changes induced by DNA damage. Mol. Cell Proteomics, 9, 457–470.

6. Deisenroth, C. and Zhang, Y. (2010) Ribosome biogenesis surveillance: probing the ribosomal protein-Mdm2-p53 pathway. Oncogene, 29, 4253–4260.

7. Rubbi, C.P. and Milner, J. (2003) Disruption of the nucleolus mediates stabilization of p53 in response to DNA damage and other stresses. EMBO J, 22, 6068–6077.

8. Woods, S.J., Hannan, K.M., Pearson, R.B. and Hannan, R.D. (2015) The nucleolus as a fundamental regulator of the p53 response and a new target for cancer therapy. Biochim. Biophys. Acta, 1849, 821–829.

9. James, A., Wang, Y., Raje, H., Rosby, R. and DiMario, P. (2014) Nucleolar stress with and without p53. Nucleus, 5.

10. Holmberg Olausson, K., Nister, M. and Lindstrom, M.S. (2012) p53-Dependent and-Independent Nucleolar Stress Responses. Cells, 1, 774–798.

11. Russo, A. and Russo, G. (2017) Ribosomal Proteins Control or Bypass p53 during Nucleolar Stress. Int J Mol Sci, 18.

12. Donati, G., Montanaro, L. and Derenzini, M. (2012) Ribosome biogenesis and control of cell proliferation: p53 is not alone. Cancer Res, 72, 1602–1607.

13. Esposito, D., Crescenzi, E., Sagar, V., Loreni, F., Russo, A. and Russo, G. (2014) Human rpL3 plays a crucial role in cell response to nucleolar stress induced by 5-FU and L-OHP. Oncotarget, 5, 11737–11751.

14. Hayden, M.S. and Ghosh, S. (2012) NF-kappaB, the first quarter-century: remarkable progress and outstanding questions. Genes Dev, 26, 203–234.

15. Karin, M. (1999) How NF-kappaB is activated: the role of the IkappaB kinase (IKK) complex. Oncogene, 18, 6867–6874.

16. DiDonato, J.A., Mercurio, F. and Karin, M. (2012) NF-kappaB and the link between inflammation and cancer. Immunol. Rev, 246, 379–400.

17. Perkins, N.D. (2012) The diverse and complex roles of NF-kappaB subunits in cancer. Nat. Rev. Cancer, 12, 121–132.

18. Pahl, H.L. (1999) Activators and target genes of Rel/NF-kappaB transcription factors. Oncogene, 18, 6853–6866.

19. Stark, L.A. and Dunlop, M.G. (2005) Nucleolar sequestration of RelA (p65) regulates NF-kappaB-driven transcription and apoptosis. Mol. Cell Biol, 25, 5985–6004.

20. Nguyen le, X.T. and Mitchell, B.S. (2013) Akt activation enhances ribosomal RNA synthesis through casein kinase II and TIF-IA. Proc. Natl. Acad. Sci. U. S. A, 110, 20681–20686.

21. Wu, S. and Tong, L. (2010) Differential signaling circuits in regulation of ultraviolet C light-induced early-and late-phase activation of NF-kappaB. Photochem. Photobiol, 86, 995–999.

22. Li, N. and Karin, M. (1998) Ionizing radiation and short wavelength UV activate NF-kappaB through two distinct mechanisms. Proc. Natl. Acad. Sci. U. S. A, 95, 13012–13017.

23. Bierhoff, H., Dundr, M., Michels, A.A. and Grummt, I. (2008) Phosphorylation by casein kinase 2 facilitates rRNA gene transcription by promoting dissociation of TIF-IA from elongating RNA polymerase I. Mol. Cell Biol, 28, 4988–4998.

24. Kato, T., Jr., Delhase, M., Hoffmann, A. and Karin, M. (2003) CK2 Is a C-Terminal IkappaB Kinase Responsible for NF-kappaB Activation during the UV Response. Mol. Cell, 12, 829–839.

25. Jiang, H.Y., Wek, S.A., McGrath, B.C., Scheuner, D., Kaufman, R.J., Cavener, D.R. and Wek, R.C. (2003) Phosphorylation of the alpha subunit of eukaryotic initiation factor 2 is required for activation of NF-kappaB in response to diverse cellular stresses. Mol. Cell Biol, 23, 5651–5663.

26. Goldstein, E.N., Owen, C.R., White, B.C. and Rafols, J.A. (1999) Ultrastructural localization of phosphorylated eIF2alpha [eIF2alpha(P)] in rat dorsal hippocampus during reperfusion. Acta Neuropathol, 98, 493–505.

27. Khandelwal, N., Simpson, J., Taylor, G., Rafique, S., Whitehouse, A., Hiscox, J. and Stark, L.A. (2011) Nucleolar NF-kappaB/RelA mediates apoptosis by causing cytoplasmic relocalization of nucleophosmin. Cell Death. Differ, 18, 1889–1903.

28. Russo, A., Pagliara, V., Albano, F., Esposito, D., Sagar, V., Loreni, F., Irace, C., Santamaria, R. and Russo, G. (2016) Regulatory role of rpL3 in cell response to nucleolar stress induced by Act D in tumor cells lacking functional p53. Cell Cycle, 15, 41–51.

29. Wan, F., Anderson, D.E., Barnitz, R.A., Snow, A., Bidere, N., Zheng, L., Hegde, V., Lam, L.T., Staudt, L.M., Levens, D. et al. (2007) Ribosomal protein S3: a KH domain subunit in NF-kappaB complexes that mediates selective gene regulation. Cell, 131, 927–939.

30. Bunz, F., Dutriaux, A., Lengauer, C., Waldman, T., Zhou, S., Brown, J.P., Sedivy, J.M., Kinzler, K.W. and Vogelstein, B. (1998) Requirement for p53 and p21 to sustain G2 arrest after DNA damage. Science, 282, 1497–1501.

31. Stark, L.A., Din, F.V.N., Zwacka, R.M. and Dunlop, M.G. (2001) Aspirin-induced activation of the NF-kB signalling pathway: A novel mechanism for aspirin-mediated apoptosis in colon cancer cells. FASEB J.

32. Voit, R. and Grummt, I. (2001) Phosphorylation of UBF at serine 388 is required for interaction with RNA polymerase I and activation of rDNA transcription. Proc. Natl. Acad. Sci. U. S. A, 98, 13631–13636.

33. Novo, S.M., Wedge, S.R. and Stark, L.A. (2017) Ex vivo treatment of patient biopsies as a novel method to assess colorectal tumour response to the MEK1/2 inhibitor, Selumetinib. Sci Rep, 7, 12020.

34. Moles, A., Sanchez, A.M., Banks, P.S., Murphy, L.B., Luli, S., Borthwick, L., Fisher, A., O’Reilly, S., van Laar, J.M., White, S.A. et al. (2013) Inhibition of RelA-Ser536 phosphorylation by a competing peptide reduces mouse liver fibrosis without blocking the innate immune response. Hepatology, 57, 817–828.

35. Drygin, D., Lin, A., Bliesath, J., Ho, C.B., O’Brien, S.E., Proffitt, C., Omori, M., Haddach, M., Schwaebe, M.K., Siddiqui-Jain, A. et al. (2011) Targeting RNA polymerase I with an oral small molecule CX-5461 inhibits ribosomal RNA synthesis and solid tumor growth. Cancer Res, 71, 1418–1430.

36. Peltonen, K., Colis, L., Liu, H., Jaamaa, S., Zhang, Z., Af, H.T., Moore, H.M., Sirajuddin, P. and Laiho, M. (2014) Small molecule BMH-compounds that inhibit RNA polymerase I and cause nucleolar stress. Mol. Cancer Ther, 13, 2537–2546.

37. Jin, R. and Zhou, W. (2016) TIF-IA: An oncogenic target of pre-ribosomal RNA synthesis. Biochim Biophys Acta, 1866, 189–196.

38. Yuan, X., Zhao, J., Zentgraf, H., Hoffmann-Rohrer, U. and Grummt, I. (2002) Multiple interactions between RNA polymerase I, TIF-IA and TAF(I) subunits regulate preinitiation complex assembly at the ribosomal gene promoter. EMBO Rep, 3, 1082–1087.

39. Mayer, C., Bierhoff, H. and Grummt, I. (2005) The nucleolus as a stress sensor: JNK2 inactivates the transcription factor TIF-IA and down-regulates rRNA synthesis. Genes Dev, 19, 933–941.

40. Mayer, C., Zhao, J., Yuan, X. and Grummt, I. (2004) mTOR-dependent activation of the transcription factor TIF-IA links rRNA synthesis to nutrient availability. Genes Dev, 18, 423–434.

41. Szymanski, J., Mayer, C., Hoffmann-Rohrer, U., Kalla, C., Grummt, I. and Weiss, M. (2009) Dynamic subcellular partitioning of the nucleolar transcription factor TIF-IA under ribotoxic stress. Biochim. Biophys. Acta, 1793, 1191–1198.

42. Chan, T.A., Morin, P.J., Vogelstein, B. and Kinzler, K.W. (1998) Mechanisms underlying nonsteroidal antiinflammatory drug-mediated apoptosis. Proc. Natl. Acad. Sci. U. S. A, 95, 681–686.

43. Charruyer, A., Grazide, S., Bezombes, C., Muller, S., Laurent, G. and Jaffrezou, J.P. (2005) UV-C light induces raft-associated acid sphingomyelinase and JNK activation and translocation independently on a nuclear signal. J. Biol. Chem, 280, 19196–19204.

44. Fillet, M., Bentires-Alj, M., Deregowski, V., Greimers, R., Gielen, J., Piette, J., Bours, V. and Merville, M.P. (2003) Mechanisms involved in exogenous C2-and C6-ceramide-induced cancer cell toxicity. Biochem. Pharmacol, 65, 1633–1642.

45. Thoms, H.C., Dunlop, M.G. and Stark, L.A. (2007) p38-mediated inactivation of cyclin D1/cyclin-dependent kinase 4 stimulates nucleolar translocation of RelA and apoptosis in colorectal cancer cells. Cancer Res, 67, 1660–1669.

46. Voit, R., Hoffmann, M. and Grummt, I. (1999) Phosphorylation by G1-specific cdk-cyclin complexes activates the nucleolar transcription factor UBF. EMBO J, 18, 1891–1899.

47. Ayrault, O., Andrique, L., Larsen, C.J. and Seite, P. (2004) Human Arf tumor suppressor specifically interacts with chromatin containing the promoter of rRNA genes. Oncogene, 23, 8097–8104.

48. Ayrault, O., Andrique, L., Fauvin, D., Eymin, B., Gazzeri, S. and Seite, P. (2006) Human tumor suppressor p14ARF negatively regulates rRNA transcription and inhibits UBF1 transcription factor phosphorylation. Oncogene, 25, 7577–7586.

49. Rocha, S., Campbell, K.J. and Perkins, N.D. (2003) p53- and Mdm2-independent repression of NF-kappa B transactivation by the ARF tumor suppressor. Mol. Cell, 12, 15–25.

50. Rothwell, P.M., Wilson, M., Elwin, C.E., Norrving, B., Algra, A., Warlow, C.P. and Meade, T.W. (2010) Long-term effect of aspirin on colorectal cancer incidence and mortality: 20-year follow-up of five randomised trials. Lancet, 376, 1741–1750.

51. Din, F.V., Theodoratou, E., Farrington, S.M., Tenesa, A., Barnetson, R.A., Cetnarskyj, R., Stark, L., Porteous, M.E., Campbell, H. and Dunlop, M.G. (2010) Effect of aspirin and NSAIDs on risk and survival from colorectal cancer. Gut, 59, 1670–1679.

52. Bywater, M.J., Poortinga, G., Sanij, E., Hein, N., Peck, A., Cullinane, C., Wall, M., Cluse, L., Drygin, D., Anderes, K. et al. (2012) Inhibition of RNA polymerase I as a therapeutic strategy to promote cancer-specific activation of p53. Cancer Cell, 22, 51–65.

53. Birbach, A., Bailey, S.T., Ghosh, S. and Schmid, J.A. (2004) Cytosolic, nuclear and nucleolar localization signals determine subcellular distribution and activity of the NF-kappaB inducing kinase NIK. J. Cell Sci, 117, 3615–3624.

54. Russo, A., Maiolino, S., Pagliara, V., Ungaro, F., Tatangelo, F., Leone, A., Scalia, G., Budillon, A., Quaglia, F. and Russo, G. (2016) Enhancement of 5-FU sensitivity by the proapoptotic rpL3 gene in p53 null colon cancer cells through combined polymer nanoparticles. Oncotarget, 7, 79670–79687.

55. Yuan, X., Zhou, Y., Casanova, E., Chai, M., Kiss, E., Grone, H.J., Schutz, G. and Grummt, I. (2005) Genetic inactivation of the transcription factor TIF-IA leads to nucleolar disruption, cell cycle arrest, and p53-mediated apoptosis. Mol. Cell, 19, 77–87.

56. Thoms, H.C., Dunlop, M.G. and Stark, L.A. (2007) CDK4 inhibitors and apoptosis: a novel mechanism requiring nucleolar targeting of RelA. Cell Cycle, 6, 1293–1297.

57. Din, F.V., Stark, L.A. and Dunlop, M.G. (2005) Aspirin-induced nuclear translocation of NFkappaB and apoptosis in colorectal cancer is independent of p53 status and DNA mismatch repair proficiency. Br. J. Cancer, 92, 1137–1143.

58. Fatyol, K. and Grummt, I. (2008) Proteasomal ATPases are associated with rDNA: the ubiquitin proteasome system plays a direct role in RNA polymerase I transcription. Biochim. Biophys. Acta, 1779, 850–859.

59. Bailly, A., Perrin, A., Bou Malhab, L.J., Pion, E., Larance, M., Nagala, M., Smith, P., O’Donohue, M.F., Gleizes, P.E., Zomerdijk, J. et al. (2016) The NEDD8 inhibitor MLN4924 increases the size of the nucleolus and activates p53 through the ribosomal-Mdm2 pathway. Oncogene, 35, 415–426.

60. Voit, R., Schafer, K. and Grummt, I. (1997) Mechanism of repression of RNA polymerase I transcription by the retinoblastoma protein. Mol. Cell Biol, 17, 4230–4237.

61. Quin, J.E., Devlin, J.R., Cameron, D., Hannan, K.M., Pearson, R.B. and Hannan, R.D. (2014) Targeting the nucleolus for cancer intervention. Biochim. Biophys. Acta, 1842, 802–816.

